# Prevention and reversion of pancreatic tumorigenesis through a differentiation-based mechanism

**DOI:** 10.1101/221986

**Authors:** Nathan M. Krah, Deanne Yugawa, Julie Straley, Christopher V. E. Wright, Raymond J. MacDonald, L. Charles Murtaugh

## Abstract

Activating mutations in *Kras* are nearly ubiquitous in human pancreatic cancer and initiate precancerous pancreatic intraepithelial neoplasia (PanINs) when induced in adult murine acinar cells. PanINs normally take months to form, but can be rapidly induced by genetic deletion of acinar cell differentiation factors such as *Ptf1a*, suggesting that loss of mature cell identity is a rate-limiting step in pancreatic tumor initiation. Using a novel genetic mouse model that allows for independent control of oncogenic *Kras* and *Ptf1a* expression, we demonstrate that maintained activity of *Ptf1a* is sufficient to eliminate Kras-driven tumorigenesis, even in the presence of tumor-promoting inflammation. Furthermore, reintroduction of *Ptf1a* into established PanINs reverts their phenotype *in vivo*. Our results suggest that reactivation of an endogenous differentiation program can prevent and reverse oncogenesis in cells harboring tumor driving mutations, thus introducing a novel paradigm for solid tumor prevention and treatment.

## INTRODUCTION

Although pancreatic ductal adenocarcinoma (PDAC) is named for its duct-like characteristics, we and others have shown that the signature driver mutation of this cancer, oncogenic *Kras*, induces tumorigenesis when introduced to exocrine acinar rather than duct cells (De La O et al., 2008; Guerra et al., 2007; Habbe et al., 2008; Ji et al., 2009; Kopp et al., 2012). Importantly, tumor initiation from *Kras*-mutant acinar cells involves a trans- or de-differentiation process, in which acinar-specific genes are downregulated and duct markers upregulated, that we refer to as “reprogramming” (De La O et al., 2008). Acinar cell-specific gene expression is normally driven by the transcription factor Ptf1a, which itself is downregulated during reprogramming to PDAC precursor lesions known as pancreatic intraepithelial neoplasia (PanIN) (De La O et al., 2008; Krah et al., 2015). Together with several cooperating transcription factors, Ptf1a is essential for maintaining mature acinar identity and restraining Kras-mediated tumorigenesis (Hoang et al., 2016; Krah et al., 2015; Shi et al., 2009; von Figura et al., 2014). How loss of Ptf1a promotes tumor development is not yet known, but could be mediated by changes in the microenvironment: acute deletion of *Ptf1a* upregulates a pro-inflammatory transcriptional program (Krah et al., 2015), and inflammation is itself known to promote PDAC (Krah and Murtaugh, 2016). Alternatively, the Ptf1a-driven transcriptional program may act cell-autonomously to suppress the effects of oncogenic *Kras;* in this scenario, the tumor-promoting effects of inflammation may be mediated via downregulation of *Ptf1a* (Molero et al., 2007). To directly test our hypothesis that enforcing acinar cell differentiation inhibits PDAC, we established an experimental system in which *Ptf1a* expression can be sustained even in the presence of oncogenic *Kras* and inflammatory injury. Our results indicate that downregulation of *Ptf1a* is essential for both basal and inflammation-driven PDAC initiation, and that reintroduction of *Ptf1a* into established tumor precursors reverts their phenotype *in vivo*.

## RESULTS

### A novel mouse model to independently control *Kras^G12D^* and *Ptf1a* expression

We established a mouse model permitting independent regulation of *Kras^G12D^* and *Ptf1a:* by tamoxifen (TM)-dependent excision of floxed STOP cassettes, *Ptf1d^CreERT^* induces expression of both *Kras^G12D^* and *rtTA*, the reverse tetracycline transactivator protein (Belteki et al., 2005) (Figure 1A-C). rtTA subsequently activates a *tetO-Ptf1a* transgene in a doxycycline (DOX)-inducible manner (Figure. 1D) (Willet et al., 2014). Expression of *rtTA* and *tetO-Ptf1a* can be monitored by their linked co-expression cassettes, *IRES-GFP* and *IRES-LacZ*, respectively. Staining for B-galactosidase (βgal) activity demonstrated acinar-specific activation of *tetO-Ptf1a* within 24 hours of DOX administration (Figure. 1E-H). Importantly, *tetO-Ptf1a* expression alone had no detectable effect on pancreas histology (Figure. 1K-L). These results indicate that both the rtTA and transgenic *Ptf1a* can be rapidly induced, specifically within Pfla-expressing cells, following TM and DOX treatment, respectively.

**Figure 1:**
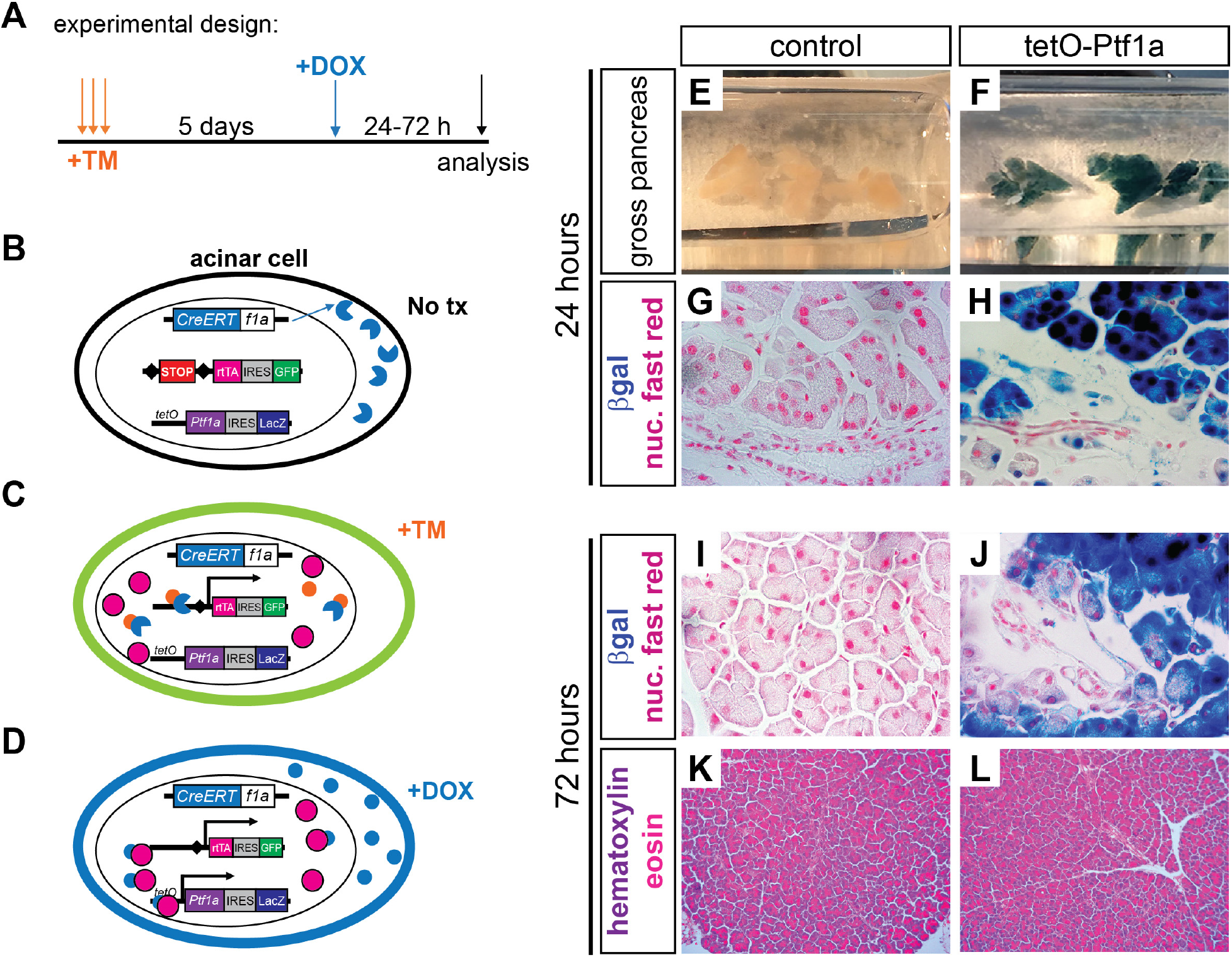
A mouse model to sustain *Ptf1a* expression in acinar cells. (**A**) Experimental schematic to induce sustained *Ptf1a* expression using alleles described below. (**B**) Prior to tamoxifen (TM) administration, CreERT fusion protein is expressed from the endogenous *Ptf1a*locus, but is sequestered to the cytoplasm. (**C**) When TM is administered, Cre-mediated recombination drives constitutive production of a reverse tetracycline transactivator protein (rtTA). Cells recombining this locus also permanently express GFP via a downstream IRES-GFP element. (**D**) Upon administration of DOX, rtTA binds tetO and drives expression of *Ptf1a*; cells activating *tetO-Ptf1a* also express β-galactosidase (βgal). (**E-F**) Wholemount βgal stained pancreata of indicated genotypes, 24 hr after DOX administration. (**G-H**) Histology of above pancreata, highlighting βgal specific to acinar cells (100x). (**I-J**) 72 hours following DOX administration, βgal remains restricted to acinar cells (100x), with no histological changes caused by *tetO-Ptf1a* expression (**K-L**, 20x).

### Sustained Ptf1a expression prevents *Kras^G12D^*-mediated pancreatic oncogenesis

To test whether sustained *Ptf1a* blocks PDAC initiation, we subjected control, *Kras^G12D^* and *Kras^G12D^*+*tetO-Ptf1a* mice (Supplementary Table 1) to high-dose TM and an 8-week (8W) chase of continuous DOX (Figure 2A; Supplementary Figure 1). All mice harbored *Ptfld^CreERT^* and *R26^LSL-rtTA^*, and exhibit uniform Cre recombination between groups (Supplementary Figure 2). After 8W, *Kras^G12D^* pancreata exhibited large areas of acinar cell loss and precancerous PanIN formation, which were greatly reduced in *Kras^GI2D^*+*tetO-Ptf1a* mice (Figure 2B-D). Regions of PDAC initiation were highlighted by expression of the ductal marker CK19 (Figure 2E-G) and mucin staining with Alcian blue (Figure 2H-J), revealing that *tetO-Ptf1a* expression dramatically reduced acinar cell transformation (Figure 2N). These results suggest that maintaining *Ptf1a* expression inhibits acinar cell transformation and prevents initiation of pancreatic tumorigenesis.

**Figure 2:**
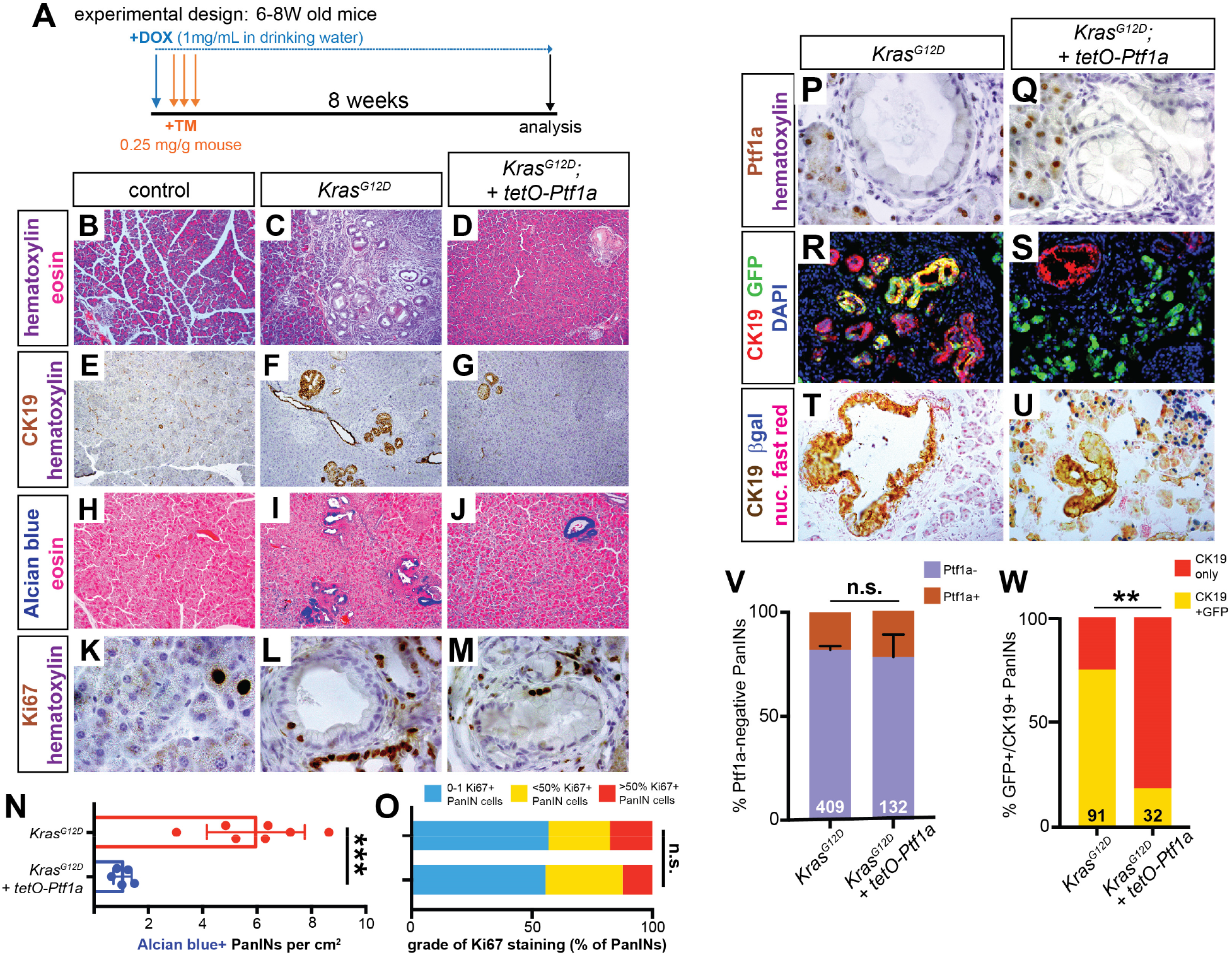
Sustained *Ptf1a* expression prevents *Kras^G12D^*-driven PanIN formation. (**A**) Mice of indicated genotypes were started on DOX (1 mg/ml in drinking water) 24 hours before TM administration (0.25 mg/g body weight, daily over 3 consecutive days). Mice were euthanized 8 weeks (8W) after the final TM treatment. (**B-D**) H&E staining of pancreata from mice of indicated genotypes 8W after TM administration (20x). Staining for CK19 (**E-G**) and Alcian blue (**H-J**) highlights reduced PanIN burden in *Kras^G12D^; tetO-Ptf1a* pancreata compared to *Kras^G12D^* alone (all images 20x). (**K-M**) Ki67 staining in PanINs of indicated mouse genotypes (100x). (N) Quantification of Alcian blue+ PanIN burden, indicating drastic reduction in lesions in *Kras^G12D^; tetO-Ptf1a* pancreata over *Kras^G12D^* alone (P<0.001). (O) Quantification of PanINs graded for proliferation based on Ki67 staining; no significant difference between genotypes (n.s.). (**P-Q**) Ptf1a staining is absent in PanINs of both genotypes, compared to adjacent acinar cells (100x). (**R-S**) Immunofluorescence for the duct marker CK19 (red), GFP (*R26^rtTA^*. green), and DAPI (blue) (20x). (**T-U**) Staining for CK19 (brown), βgal *(tetO-Ptf1a*, blue) and nuclear fast red (40x). (**V**) Proportion of PanINs that are completely Ptf1a negative vs. those that harbor one or more Ptf1a-positive cell(s). (**W**) Proportion of CK19+/GFP+ dual-positive PanINs (Fisher’s exact test, *p*<0.01). Numbers on bar graphs indicate total lesions scored, across 3-5 mice per group. \*\**p*<0.01, \*\*\**p*<0.001.

As rare PanINs were still generated in *Kras^G12D^* + *tetO-Ptf1a* pancreata, we compared these lesions to PanINs induced by *Kras^G12D^* alone. Neither the amount of proliferation (Figure 2K-M, O) nor the number of Ptf1a+ cells within PanINs differed between genotypes (Figure 2P-Q, V), suggesting that PanINs were similar in both groups. We therefore postulated that PanINs in *Kras^G12D^* +*tetO-Ptf1a* mice were “escapers,” recombining *Kras^G12D^* to initiate tumorigenesis, but not *R26^LSL-rtTA^*, which is required for sustained *Ptf1a* expression. To test this, we determined the frequency of CK19+ PanINs containing GFP+ cells. Most *Kras^G12D^* PanINs co-expressed CK19 and GFP, indicating derivation from acinar cells recombining both the *Kras^G12D^* and *R26^LSL-rtTA^* loci (Figure 2R, W). In contrast, almost all *Kras^G12D^* + *tetO-Ptf1a* PanINs were broadly GFP-negative (Figure 2S, W), indicating that they failed to recombine *R26^LSL-rtTA^*. Thus, incomplete Cre-based recombination explains residual PanIN formation in *Kras^G12D^* + *tetO-Ptf1a*pancreata. βgal staining further confirmed that *Kras^G12D^* + *tetO-Ptf1a* PanINs contained only very rare cells expressing *tetO-Ptf1a* (Figure 2T-U). Taken together, these results indicate that *Ptf1a* expression is sufficient to prevent PDAC initiation.

### Pancreatitis is insufficient to overcome Pfla-mediated tumor suppression

As noted above, deletion of *Ptf1a* enhances pancreatic inflammation (Krah et al., 2015), which could mediate its pro-tumorigenic consequences. To determine if inflammation could bypass tumor suppression by *Ptf1a*, we activated *tetO-Ptf1a* in *Kras^G12D^* mice that were also subjected to caerulein-induced pancreatitis, a model of increased PDAC risk (Fig 3A) (Guerra et al., 2007; Lowenfels et al., 1993). Pancreata of control mice were fully recovered at 3 weeks following induction of pancreatitis, while robust PanIN formation was seen in *Kras^G12D^* pancreata, as previously reported (Fig. 3B-C) (Guerra et al., 2007; Kopp et al., 2012). In contrast, *Kras^G12D^* + *tetO-Ptf1a* pancreata exhibited reduced PanIN formation and lacked the fibro-inflammatory stroma characteristic of caerulein-treated *Kras^G12D^* mice (Fig. 3D). CK19 and Alcian blue analysis confirmed the dramatic reduction in PanIN burden in *Kras^G12D^* + *tetO-Ptf1a* pancreata (Fig. 3E-J, Q). The absence of anti-GFP and anti-Ptf1a staining again indicated that residual PanINs in these mice arose from Ptf1a-negative escaper cells (Fig. 3K-P, R). Thus, *Ptf1a* is both necessary and sufficient to block PDAC initiation, even in the presence of tumor-promoting inflammation (Krah et al., 2015). These results additionally indicate that downregulation of Ptf1a is a necessary cell-autonomous event for PanIN initiation, as only cell incapable of sustaining Ptf1a expression (i.e. non-rtTA-expressing, GFP-negative cells) give rise to PanINs.

**Figure 3:**
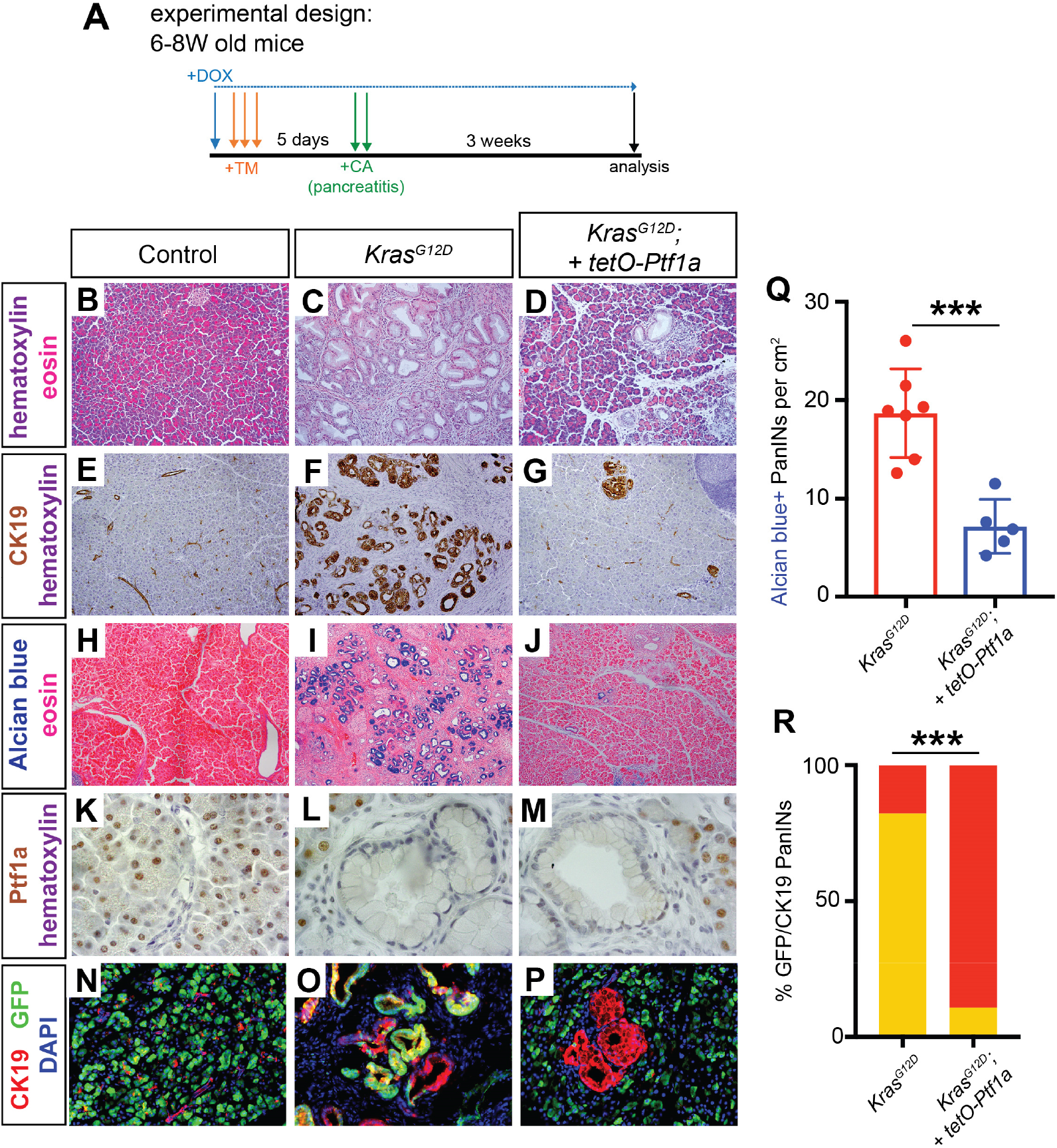
Maintenance of acinar identity inhibits inflammation-induced PanIN formation. (**A**) Mice of indicated genotypes were started on DOX (1 mg/ml in drinking water) 24 hours before TM administration (0.25 mg/g body weight, daily over 3 consecutive days). Five days following the final TM dose, mice were administered six hourly injections of 0.1 μg/g caerulein on two consecutive days. Mice were euthanized for pancreatic histology at three weeks following the final caerulein injection. (**B-D**) H&E staining of pancreata from mice of indicated genotypes 3 weeks after caerulein-induced pancreatitis (20x). (**E-G**) IHC for CK19 (20x) and (**H-J**) Alcian blue staining (10x), highlighting PanIN formation. (**K-M**) IHC for Ptf1a (100x), showing lack of nuclear Ptf1a in normal ducts of control mice as well as in PanINs, compared to adjacent acinar cells. (**N-P**) Immunofluorescence for the duct marker CK19 (red), GFP (*R26^rtTA^*. green) and DAPI (blue), highlighting CK19+/GFP-negative escaper PanIN cells in *Kras^G12D^; tetO-Ptf1a* pancreata (20x). (**Q**) Quantification of the genotype-dependent PanIN burden, indicating significant reduction in *Kras^G12D^; tetO-Ptf1a* pancreata (unpaired t-test, P<0.01). (R) Proportion of CK19+/GFP+ dual-positive PanINs (Fisher’s exact test, P<0.001).

### Re-expression of *Ptf1a* reverses pancreatic transformation *in vivo*

The therapeutic potential of these findings would be greatest if *Ptf1a* activation could redirect the cell fate of already-established PDAC precursors, preventing tumor progression and reverting transformed cells to a stable differentiated state. To address this, we allowed PanINs to form in DOX-untreated *Kras^G12D^* and *Kras^G12D^* + *tetO-Ptf1a* mice for 8W (**DOX-off**), followed by DOX-induction of Ptf1a for 3W (**DOX-on**). Whereas large PanINs were present in *Kras^G12D^* pancreata, lesions of *Kras^G12D^*+*tetO-Ptf1a* mice appeared smaller and contained misplaced Ptf1a+ cells (Supplementary Figure 3). To specifically follow the fate of Pfla-reactivating cells within established lesions, we analyzed GFP (rtTA) and βgal *(tetO-Ptf1a)* production at multiple timepoints before and after DOX (Figure 4). Most PanINs that formed at 8W DOX-off, or those analyzed after 24 hrs DOX-on after 8W DOX-off, contained abundant GFP+ cells (Figure 4A, J), i.e., they expressed rtTA. By contrast, 3W and 6W DOX-on treatment induced a progressive exclusion of GFP+ cells from PanINs (Figure 4B-C, J). At 3W DOX-on, we observed a striking formation of hybrid structures in which clustered GFP+ cells appeared tightly connected to CK19+ lesions (Figure 4B’, K). In contrast to *Kras^G12D^* pancreata where amylase+ cells were excluded from PanINs, the emerging GFP+ cells in *Kras^G12D^*+ tetO-Ptf1a pancreata were amylase+, suggesting re-differentiation into acini (Figure 4D-E, white arrows). These results were corroborated by the LacZ (*tetO-Ptf1a*) reporter: while DOX-off mice had no βgal+ pancreatic cells, 24 hr DOX-on induced robust βgal expression in a subset of PanINs, which were then absent from lesions after 3-6W DOX-on (Figure 4F-I, L). Together, these data suggest that *Ptf1a* re-expression reversed transformation of pre-cancerous cells *in vivo*.

**Figure 4:**
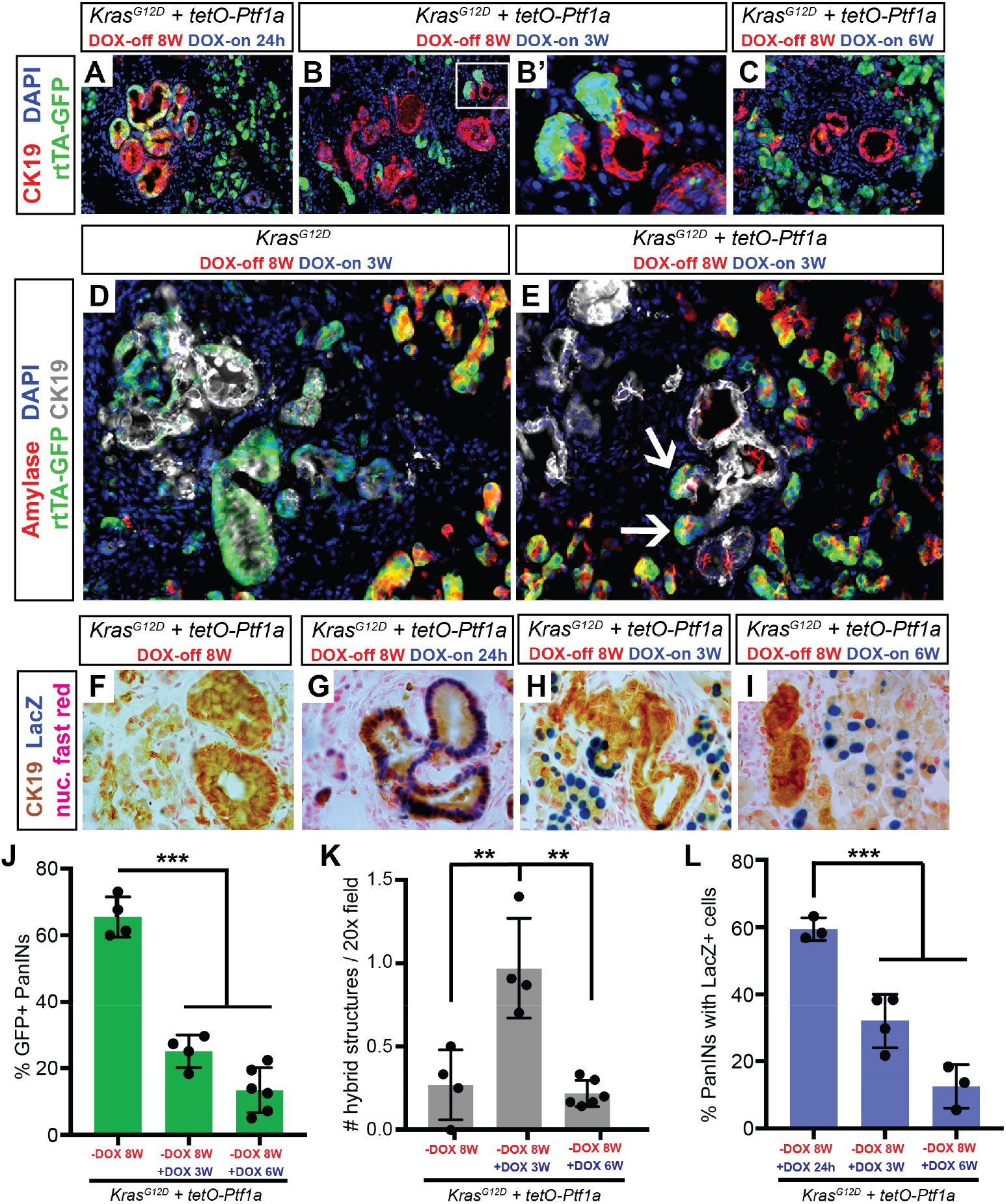
Re-activation of *Ptf1a* in PanINs reverses premalignant cell phenotypes. (**A-C**) Immunofluorescence for CK19 (red), GFP (*R26^rtTA^*, green), and DAPI (blue) in *Kras^G12D^+ tetO-Ptf1a* pancreata following indicated treatments (20x). (B’) Enlarged image highlighting GFP+ acinar cells emerging from CK19+ lesions. (**D-E**) Immunofluorescence for amylase (red), GFP (*R26^rtTA^*, green), CK19 (white), and DAPI (blue) in *Kras^G12D^* and *Kras^G12D^*+*tetO-Ptf1a*pancreata following indicated treatment (20x). (E) Arrows indicate Amylase/GFP-dual positive cells emerging from CK19+ lesions. (**F-I**) Staining for CK19 (brown), βgal (*tetO-Ptf1a*, blue) and nuclear fast red in pancreata of indicated treatment groups (100x). (**J-L**) Quantification of proportion of GFP+ PanINs (**J**), number of PanIN/acinar hybrid structures per field (**K**), and proportion of βgal+ PanINs (L) for mice of indicated treatment groups. ** *p*<0.01, *** *p*<0.001, as determined by ANOVA.

## DISCUSSION

Pancreatic cancer evolves through premalignant stages lasting a decade or more, providing a window for prevention and reversion of this deadly disease with the development of improved tools for detection and management (Hruban et al., 2000; Yachida et al., 2010). Several recent mouse modeling studies have shown the importance of acinar cell differentiation factors, including *Ptf1a*, in restraining PDAC initiation (Flandez et al., 2014; Krah et al., 2015; Roy et al., 2016; Shi et al., 2009; von Figura et al., 2014). Consistent with these findings, human genome-wide association studies have identified polymorphisms in *NR5A2* and *PDX1*,transcriptional partners and regulators of *Ptf1a*, which increase the risk of PDAC (Petersen et al., 2010; Wolpin et al., 2014). *Ptf1a* itself is downregulated in human PanINs, and required in mice to maintain homeostasis, restrain pancreatic inflammation, and suppress RAS-related gene signatures (Hoang et al., 2016; Krah et al., 2015). Together, these studies suggest that maintaining acinar cell differentiation could be an attractive therapeutic approach to limit pancreatic tumor initiation and progression (Murtaugh, 2014; Rooman and Real, 2012).

Such an approach is supported by the findings reported here, that sustained *Ptf1a* expression can prevent and reverse the early stages of pancreatic tumorigenesis. Our results indicate that downregulation of *Ptf1a* is essential for PanIN formation and maintenance, as only Ptf1a-negative “escaper” cells contribute to PanINs in *Kras^G12D^*+*tetO-Ptf1a* pancreata, in two different experimental schemas of PDAC initiation (Figures 2 and 3). Additionally, transgenic reintroduction of *Ptf1a* into established lesions reverts their phenotype to amylase-positive acinar cells, providing the first functional evidence that differentiation can override *Kras^G12D^*-mediated oncogenesis (Figure 4). These findings are consistent with a previous study, which suggested that PanIN cells could revert to normal acini following the silencing of *Kras^Gl21D^* (Collins et al., 2012). As previous work from our labs suggests that *Ptf1a* suppresses RAS-dependency genes (Krah et al., 2015), we speculate that activation of *Ptf1a* indirectly inhibits transformation by inhibiting expression of genes required for oncogenic *Kras* function, while also promoting tissue reorganization, maintenance, and homeostasis (Hoang et al., 2016). Importantly, while *Kras* activation is an effectively irreversible genetic alteration in PDAC, downregulation of *Ptf1a* occurs at the transcriptional level, and is in principle reversible. Strategies aimed at restoring *Ptf1a* expression should complement efforts to develop inhibitors of KRAS itself and its downstream effectors (Collins and Pasca di Magliano, 2013; McCormick, 2015).

Chemoprevention strategies are under active investigation in PDAC (Miller et al., 2016), and our findings suggest that maintenance of *Ptf1a* and other acinar cell differentiation factors will be pivotal to their success, and could enable PDAC prevention even in high-risk individuals with chronic inflammation. Our results highlight the capacity of an epigenetic differentiation program to overcome the genetic alterations driving tumorigenesis, which may apply beyond PDAC to other solid tumors in which initiation involves alterations of differentiation state (Krah and Murtaugh, 2016; Roy and Hebrok, 2015).

## AUTHOR CONTRIBUTIONS

N.M.K., R.M.J. and L.C.M. designed the research study. N.M.K, D.Y., J.S, and L.C.M acquired and analyzed the data. C.V.E.W. provided pivotal reagents. The manuscript was written by N.M.K and L.C.M. with input from R.M.J. and C.V.E.W.

## ACKNOWLEDGMENTS

We are grateful to members of our laboratories as well as Howard Crawford, Gabrielle Kardon and Matthew Firpo for helpful input. This work was supported by the National Institutes of Health, through the following grants: F30-CA192819 (N.M.K.), R01-DK061220 and R01-CA194941 (L.C.M. and R.J.M.); U01-DK089570 (C.V.E.W.). We declare no conflicts of interest.

## METHODS

Further information and requests for resources and reagents should be directed to and will be fulfilled by the corresponding author, L. Charles Murtaugh.

### EXPERIMENTAL MODEL AND SUBJECT DETAILS

#### Mice

Experimental mouse alleles have been utilized in previous publications within the pancreatic research community: *Ptf1a^CreERT^* (*Ptf1a^tm2(CreER1)CVW^*) (Kopinke et al., 2012; Kopp et al., 2012; Krah et al., 2015), *Kras^LSL-GI2D^* (*Kras^tm4ty^*]) (Hingorani et al., 2003), *Rosa26^rtTA-IRES-EGFP^* (Belteki et al., 2005; Collins et al., 2012), and *tetO-Ptf1a* (Willet et al., 2014). To induce Cre-mediated recombination, tamoxifen (Cayman Chemical, Ann Arbor, MI) in corn oil was administered via oral gavage at 0.25 mg/g body weight on three consecutive days. To induce *tetO-Ptf1a* expression, all experimental mice were administered 1 mg/mL of Doxycycline (Research Products International, Mt. Prospect, IL) with 1% D-sucrose (Fisher Scientific) in their drinking water. Water bottles containing Doxycycline were protected from light and refreshed two times per week. All mice utilized in this study were between 6 and 12 weeks of age at the beginning of experiments. No mice with health status reports, as determined by an in-house veterinarian, were utilized in any analysis. For each experiment and experimental group, we utilized approximately identical number of male and female mice. Previous studies have <u>not</u> identified a gender-dependent PanIN burden in any mouse model of PDAC initiation or progression. Furthermore, human PDAC affects men and women with similar incidence, leading to ~7% of cancer related deaths in both genders (Siegel et al., 2016).

All experiments involving mice were performed according to institutional IACUC and NIH guidelines.

#### Tissue processing and histology

Pancreata were dissected into ice cold PBS, separated into multiple parts and processed for frozen and paraffin sections, as previously described (De La O et al., 2008; Krah et al., 2015). For paraffin sectioning, tissues were fixed in zinc-buffered formalin (Z-fix; Anatech, Battle Creek, MI) at room temperature overnight, followed by processing (dehydration in ethanol washes) into Paraplast-Plus (McCormick Scientific). Frozen specimens were fixed for 1-2 hr in 4% paraformaldehyde in 1x PBS on ice, followed by processing into Tissue-Tek O.C.T. compound (Fisher Healthcare). Paraffin and frozen sections were 6-8-microns with ≥100 μm spacing between individual sections, all placed on a single slide.

IHC and immunofluorescence followed established protocols (De La O et al., 2008; Krah et al., 2015) and included high temperature antigen retrieval (Vector Unmasking Solution; Vector Laboratories, Burlingame, CA), prior to staining all paraffin sections. Primary antibodies are listed in the Key Resource Table. Secondary antibodies, raised in donkey (Jackson Immunoresearch, West Grove, PA) were diluted 1:250 in blocking solution. Vectastain reagents and diaminobenzidine (DAB) substrate (Vector Laboratories) were used for all IHC experiments (See Key Resource Table). Immunofluorescence sections were counterstained with DAPI and mounted in Fluoromount-G (Southern Biotech), and photographed on an Olympus IX71 microscope, using MicroSuite software (Olympus America, Waltham, MA). Images were processed in Adobe Photoshop, with exposure times and adjustments identical between genotypes and treatment groups.

For Alcian blue staining, paraffin sections were pre-incubated 15 min in 3% acetic acid, stained 10 min in 1% Alcian blue in 3% acetic acid), and washed extensively in 3% acetic acid and dH2O. Following staining, all slides were washed in 0.5% acetic acid, dehydrated and equilibrated into xylene, and mounted with Permount.

### METHOD DETAILS

#### Quantifications

##### PanIN scoring

To measure the number of PanINs per pancreas, the entire surface area of each Alcian blue/eosin-stained section was photographed at 4x original magnification, followed by photo-merging in Adobe Photoshop. The surface area was measured using ImageJ software (NIH). Alcian blue+ PanINs were counted manually under the microscope and marked on composite images in Adobe Photoshop. PanIN burden was calculated as total number of Alcian blue+ lesions per cm^2^. As previously described, metaplastic lesions that did not stain with Alcian blue were not counted (Krah et al., 2015). To avoid double-counting tortuous lesions that could occupy multiple regions in 3-D space, no more than one lesion was scored within an anatomically distinct pancreatic lobule (De La O et al., 2008; Krah et al., 2015).

##### Quantification of immunofluorescence images

To quantify the *R26^rtTA^* recombination frequency, we imaged 10-12 randomly selected 20x fields per specimen (across multiple sections). Using ImageJ (NIH), cell co-expressing GFP with the acinar differentiation marker, Amylase, were detected by additive image overlay with DAPI and anti-GFP, and counted using the Analyze Particles function as described previously (Keefe et al., 2012; Krah et al., 2015). To ensure counting accuracy, random images were manually spot-checked, using Adobe Photoshop. All calculations were performed in Microsoft Excel and results graphed as individuals with error bars representing the standard deviation. The p-values were determined by two-tailed, unpaired t-test in Graphpad Prism 7.

To quantify the number of GFP-positive PanINs, 10 randomly selected fields were imaged per mouse. Each PanIN was manually scored according to the number of GFP+/CK19+ cells present. If more than two cells (or regions) co-expressed CK19 and GFP, the PanIN was considered GFP-positive.

##### Quantification of histological images

To quantify cell proliferation in PanINs, each lesion was scored according to its number of Ki67+ nuclei. Each lesion was categorized as low-(0-1 Ki67+ nuclei), mid-(<50% Ki67+ nuclei), or high-grade (>50% Ki67+ nuclei). At least 10 fields with PanINs were counted per animal (n=3 mice per genotype). To quantify the number of Ptf1a-negative vs. positive PanINs, as many lesions as possible were imaged from Ptf1a-stained *Kras^G12D^* and *Kras^G12D^* + *tetO-Ptf1a* pancreata (n=5 mice per genotype). If two or more cells contained Ptf1a+ nuclei, the PanIN was considered Ptf1a-positive. The number of βgal+ PanINs was determined by counting βgal+ cells per lesion; if the lesion contained 2 or more βgal (tetO-Ptf1a)-positive cells, the lesion was considered positive.

##### Caerulein treatment

Acute pancreatitis was induced by i.p. injection of caerulein (Bachem, Torrance, CA), 0.1 μg/g in filter sterilized saline, six times (every 60 minutes) daily over two consecutive days, as previously described (Keefe et al., 2012; Kopp et al., 2012; Krah et al., 2015). Controls were injected with an equal volume of sterile saline. Pancreata from all caerulein-treated mice were harvested three weeks following final injection and processed as described above.

### QUANTIFICATION AND STATISTICAL ANALYSIS

#### Statistics

All statistics were performed in Graphpad Prism 7. A two-tailed t-test was used to calculate p-values for PanIN burden and percentage of Ptf1a-negative PanINs (Figure 2). A Fisher exact test was used to calculate *p*-values for nominal data, such as relative frequencies of GFP/CK19 PanINs (Figure 2 and Figure 3). Where multiple groups were compared against one another (Figure 4), *p*-values were determined by ANOVA. All graphs, regardless of the statistical test used, show the mean ± the standard deviation. Statistical significance was defined as a *P*-value of <0.05 for the indicated analysis, as determined by Graphpad Prism 7 software.

### KEY RESOURCE TABLE

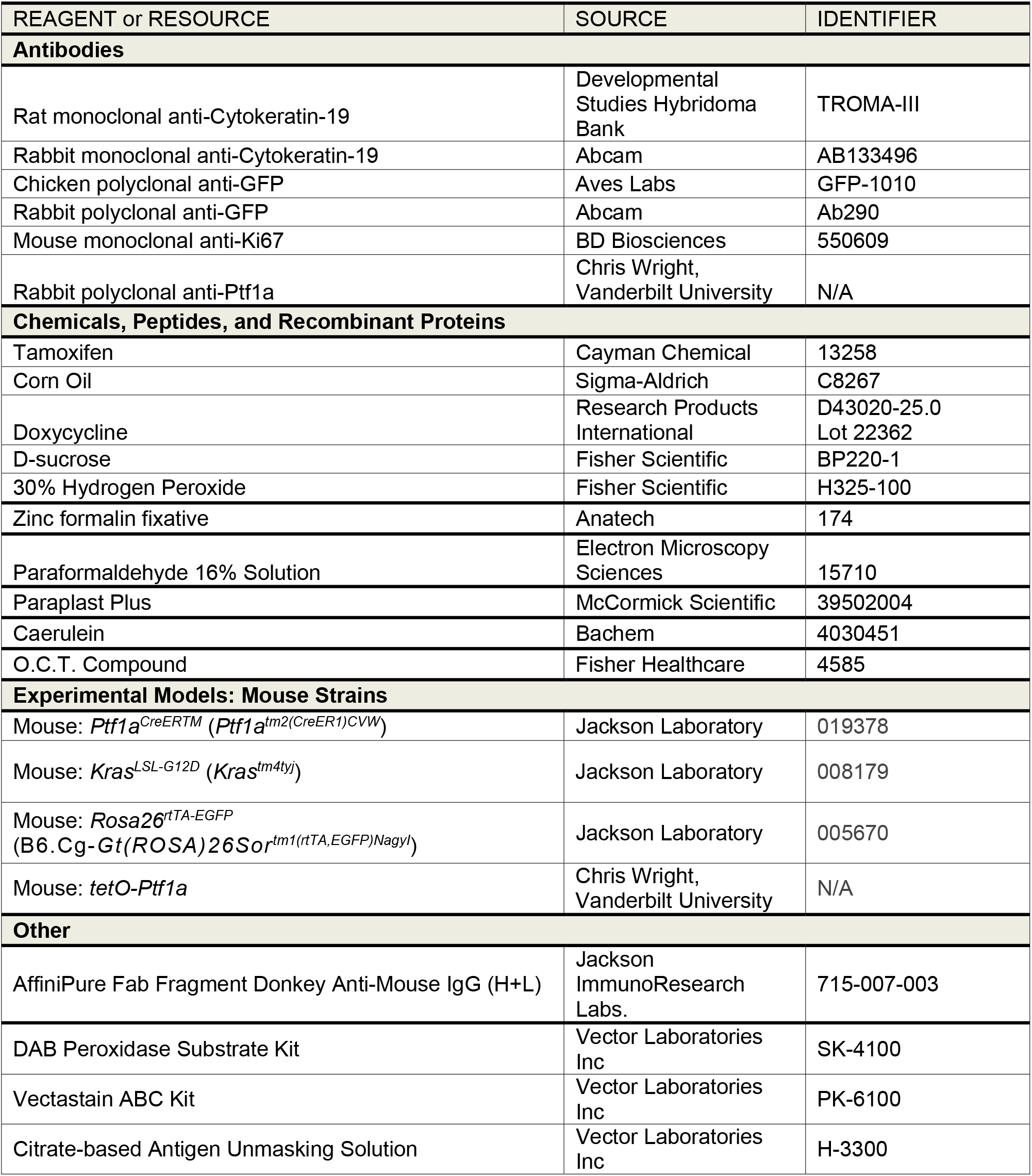

**Supplementary Table 1.**
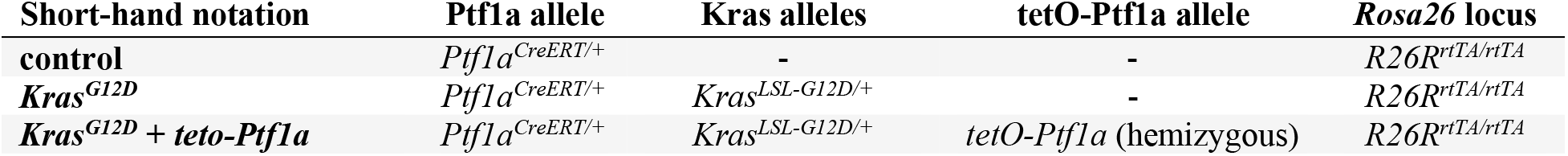
Nomenclature of mouse mutants used in this study

**Supplementary Table 2.**
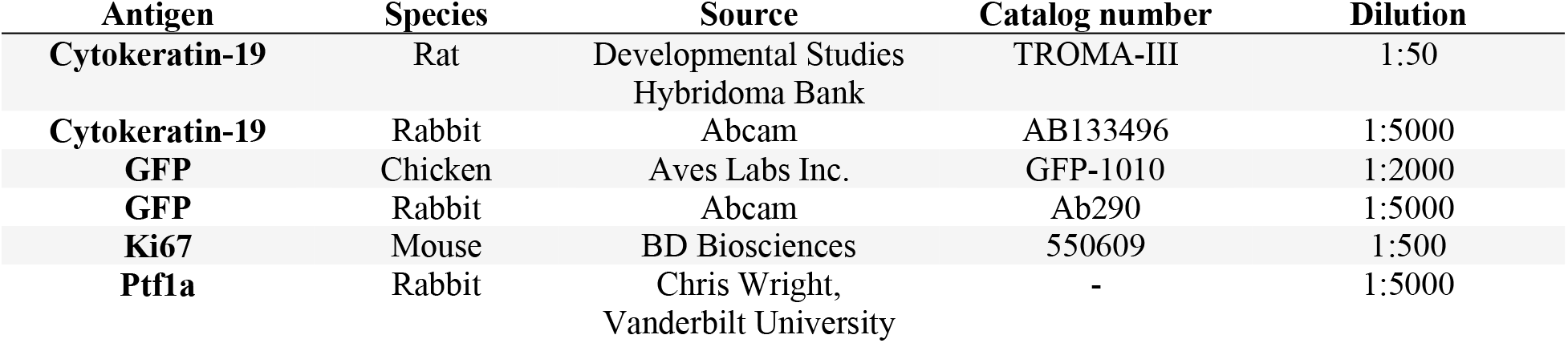
Primary antibodies utilized in this study

## SUPPLEMENTARY FIGURE

**supplementary Figure 1:**
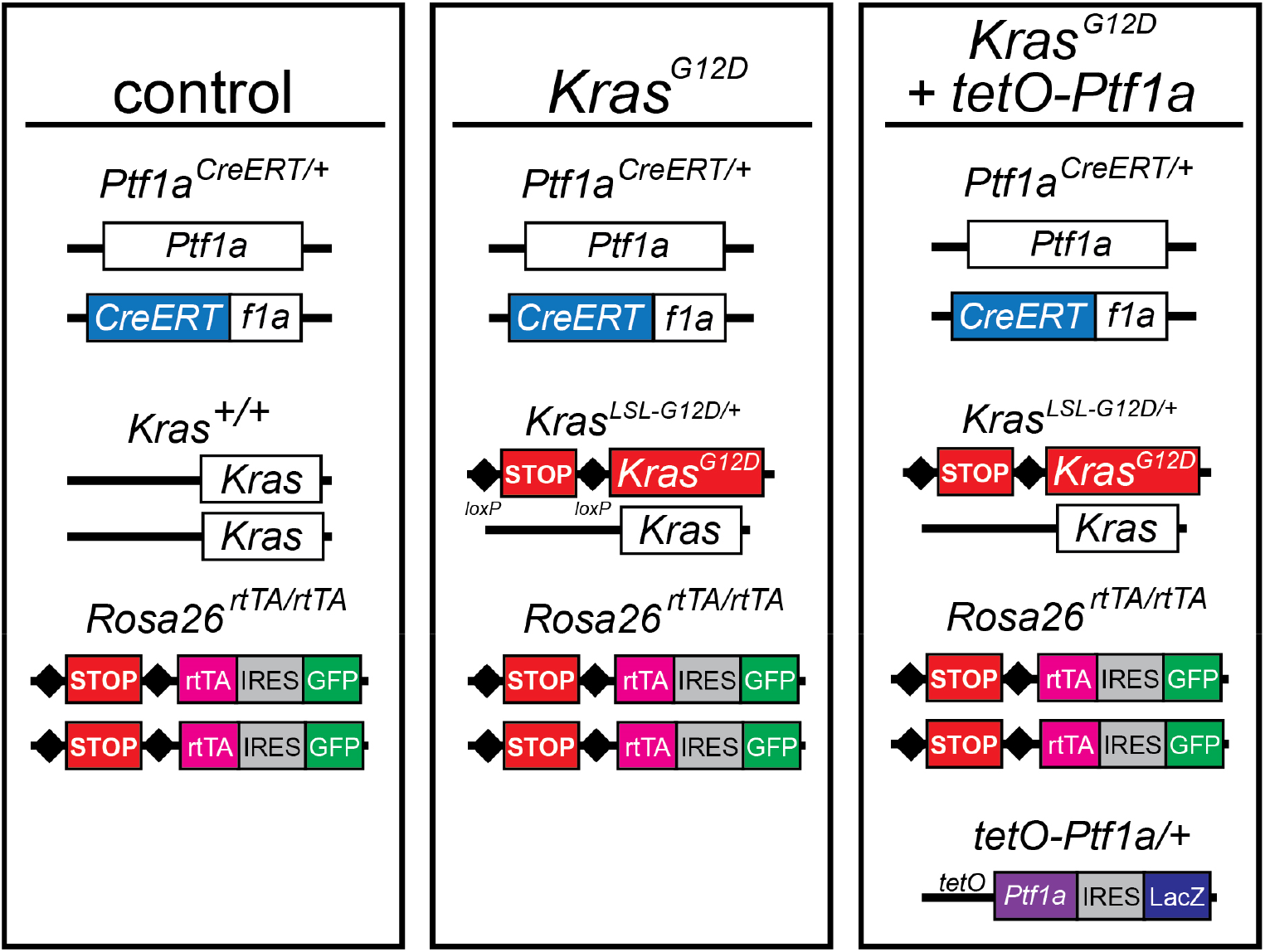
Mouse alleles utilized in this study. Schematic representations of the alleles present in the genotypes referred to, in shorthand, as Control, *Kras^G12D^*, and *Kras^G12D^* + *tetO-Ptf1a*. Function of the system is described in Figure 1.

**supplementary Figure 2:**
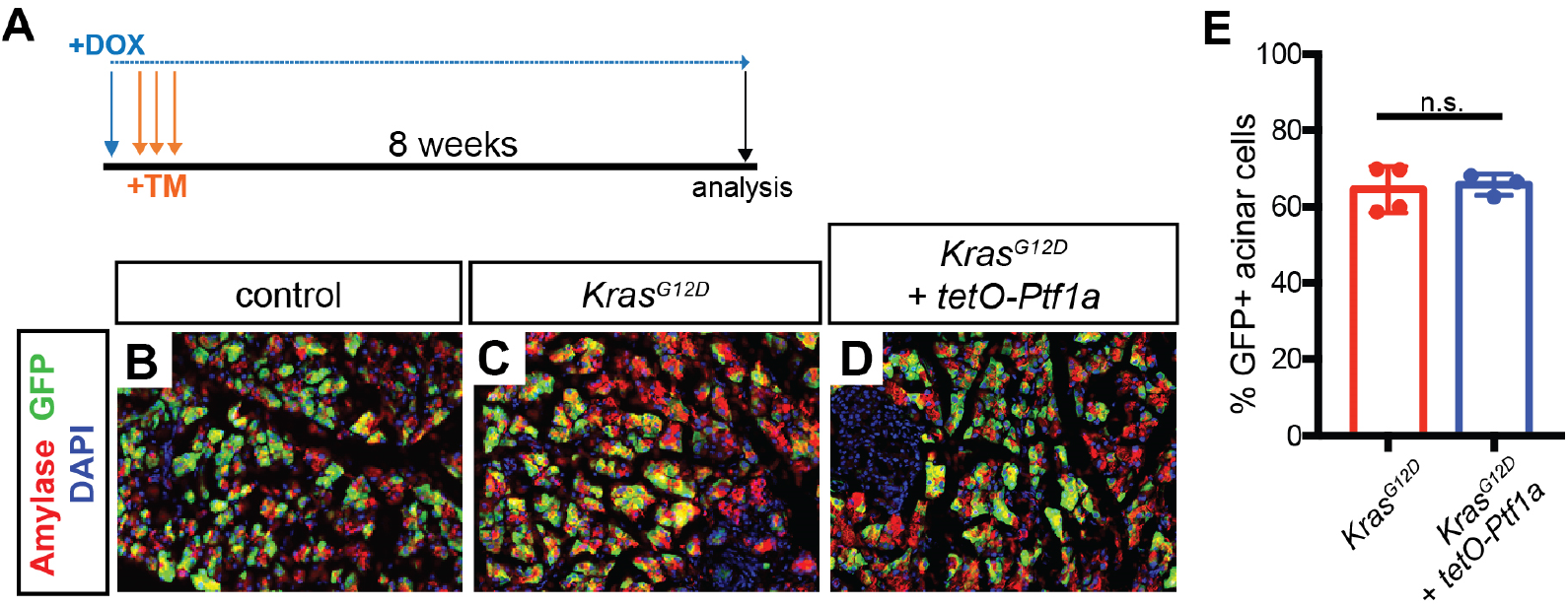
*Ptf1a^CreERT^* recombination efficiency following tamoxifen treatment. (A) 6-8-week-old mice were administered tamoxifen on three consecutive days (3 × 0.25 mg/g mouse) while DOX was present in the drinking water (1 mg/ml), and pancreata were harvested after an 8-week chase. (**B-D**) Immunofluorescence for amylase (red) and GFP (green), reporting recombination of *R26^rTA^*, on pancreata of indicated genotypes (20x). For all mice, only histologically normal areas were imaged in an effort to provide an accurate quantification of Cre-mediated recombination. (**E**) The proportion of GFP expression among amylase+ acinar cells were compared between *Kras^G12D^*, and *Kras^G12D^* + *tetO-Ptf1a* genotypes, with no significant difference found.

**supplementary Figure 3:**
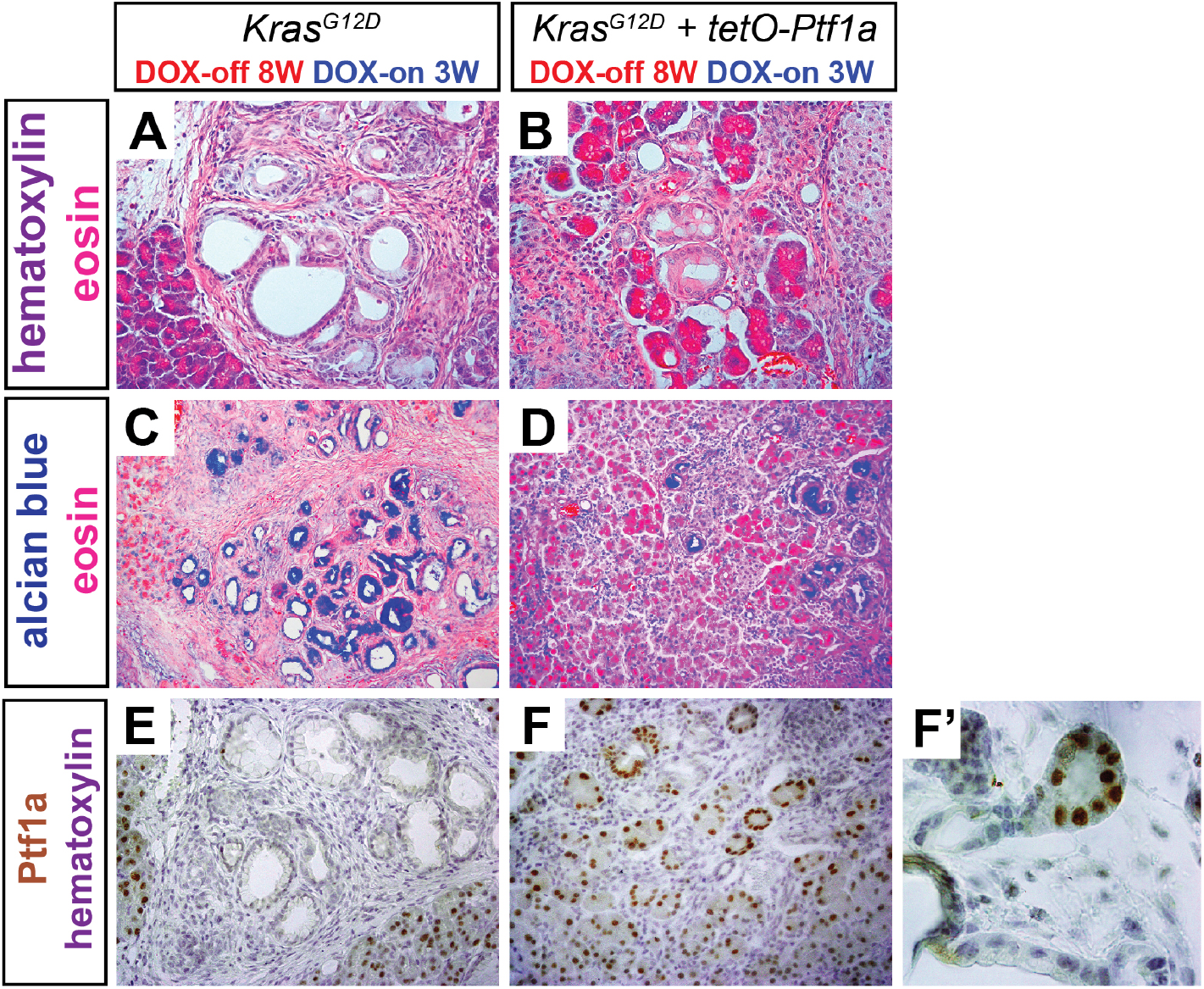
*Ptf1a* re-expression in PanINs leads to the emergence of lesion-localized acinar cells in *Kras^G12D^*+*tetO-Ptf1a* pancreata. (**A-B**) H&E (40x) and (**C-D**) Alcian blue (20x) staining of pancreata from mice of indicated genotypes and treatments. Eosinophilic primitive acinar cells are observed trapped within PanIN lesions of *Ptf1a* re-expressing mice. (**E-F**) Immunohistochemistry for Ptf1a, showing absence of Ptf1a in PanINs from *Kras^G12D^*pancreata and wide Ptf1a expression in duct-like structures in *Kras^G12D^*+*tetO-Ptf1a* pancreata (40x). (F’) Ptf1a+ acinar-like cluster emerging from a ductal structure.

## REFERENCES

Belteki, G., Haigh, J., Kabacs, N., Haigh, K., Sison, K., Costantini, F., Whitsett, J., Quaggin, S.E., and Nagy, A. (2005). Conditional and inducible transgene expression in mice through the combinatorial use of Cre-mediated recombination and tetracycline induction. Nucleic Acids Res 33, e51.

Collins, M.A., Bednar, F., Zhang, Y., Brisset, J.C., Galban, S., Galban, C.J., Rakshit, S., Flannagan, K.S., Adsay, N.V., and Pasca di Magliano, M. (2012). Oncogenic Kras is required for both the initiation and maintenance of pancreatic cancer in mice. J Clin Invest 122, 639–653.

Collins, M.A., and Pasca di Magliano, M. (2013). Kras as a key oncogene and therapeutic target in pancreatic cancer. Front Physiol 4, 407.

De La O, J.P., Emerson, L.L., Goodman, J.L., Froebe, S.C., Illum, B.E., Curtis, A.B., and Murtaugh, L.C. (2008). Notch and Kras reprogram pancreatic acinar cells to ductal intraepithelial neoplasia. Proc Natl Acad Sci U S A 105, 18907–18912.

Flandez, M., Cendrowski, J., Canamero, M., Salas, A., del Pozo, N., Schoonjans, K., and Real, F.X. (2014). Nr5a2 heterozygosity sensitises to, and cooperates with, inflammation in KRas(G12V)-driven pancreatic tumourigenesis. Gut 63, 647–655.

Guerra, C., Schuhmacher, A.J., Canamero, M., Grippo, P.J., Verdaguer, L., Perez-Gallego, L., Dubus, P., Sandgren, E.P., and Barbacid, M. (2007). Chronic pancreatitis is essential for induction of pancreatic ductal adenocarcinoma by K-Ras oncogenes in adult mice. Cancer Cell 11, 291–302.

Habbe, N., Shi, G., Meguid, R.A., Fendrich, V., Esni, F., Chen, H., Feldmann, G., Stoffers, D.A., Konieczny, S.F., Leach, S.D., et al. (2008). Spontaneous induction of murine pancreatic intraepithelial neoplasia (mPanIN) by acinar cell targeting of oncogenic Kras in adult mice. Proc Natl Acad Sci U S A 105, 18913–18918.

Hingorani, S.R., Petricoin, E.F., Maitra, A., Rajapakse, V., King, C., Jacobetz, M.A., Ross, S., Conrads, T.P., Veenstra, T.D., Hitt, B.A., et al. (2003). Preinvasive and invasive ductal pancreatic cancer and its early detection in the mouse. Cancer Cell 4, 437–450.

Hoang, C.Q., Hale, M.A., Azevedo-Pouly, A.C., Elsasser, H.P., Deering, T.G., Willet, S.G., Pan, F.C., Magnuson, M.A., Wright, C.V., Swift, G.H., et al. (2016). Transcriptional Maintenance of Pancreatic Acinar Identity, Differentiation, and Homeostasis by PTF1A. Mol Cell Biol 36, 3033–3047.

Hruban, R.H., Goggins, M., Parsons, J., and Kern, S.E. (2000). Progression model for pancreatic cancer. Clin Cancer Res 6, 2969–2972.

Ji, B., Tsou, L., Wang, H., Gaiser, S., Chang, D.Z., Daniluk, J., Bi, Y., Grote, T., Longnecker, D.S., and Logsdon, C.D. (2009). Ras activity levels control the development of pancreatic diseases. Gastroenterology 137, 1072–1082, 1082 e1071-1076.

Keefe, M.D., Wang, H., De La, O.J., Khan, A., Firpo, M.A., and Murtaugh, L.C. (2012). betacatenin is selectively required for the expansion and regeneration of mature pancreatic acinar cells in mice. Dis Model Mech 5, 503–514.

Kopinke, D., Brailsford, M., Pan, F.C., Magnuson, M.A., Wright, C.V., and Murtaugh, L.C. (2012). Ongoing Notch signaling maintains phenotypic fidelity in the adult exocrine pancreas. Dev Biol 362, 57–64.

Kopp, J.L., von Figura, G., Mayes, E., Liu, F.F., Dubois, C.L., Morris, J.P.T., Pan, F.C., Akiyama, H., Wright, C.V., Jensen, K., et al. (2012). Identification of Sox9-dependent acinar-to-ductal reprogramming as the principal mechanism for initiation of pancreatic ductal adenocarcinoma. Cancer Cell 22, 737–750.

Krah, N.M., De La, O.J., Swift, G.H., Hoang, C.Q., Willet, S.G., Chen Pan, F., Cash, G.M., Bronner, M.P., Wright, C.V., MacDonald, R.J., et al. (2015). The acinar differentiation determinant PTF1A inhibits initiation of pancreatic ductal adenocarcinoma. eLife 4.

Krah, N.M., and Murtaugh, L.C. (2016). Differentiation and Inflammation: ‘Best Enemies’ in Gastrointestinal Carcinogenesis. Trends Cancer 2, 723–735.

Lowenfels, A.B., Maisonneuve, P., Cavallini, G., Ammann, R.W., Lankisch, P.G., Andersen, J.R., Dimagno, E.P., Andren-Sandberg, A., and Domellof, L. (1993). Pancreatitis and the risk of pancreatic cancer. International Pancreatitis Study Group. N Engl J Med 328, 1433–1437.

McCormick, F. (2015). KRAS as a Therapeutic Target. Clin Cancer Res 21, 1797–1801.

Miller, M.S., Allen, P., Brentnall, T.A., Goggins, M., Hruban, R.H., Petersen, G.M., Rao, C.V., Whitcomb, D.C., Brand, R.E., Chari, S.T., et al. (2016). Pancreatic Cancer Chemoprevention Translational Workshop: Meeting Report. Pancreas 45, 1080–1091.

Molero, X., Adell, T., Skoudy, A., Padilla, M.A., Gomez, J.A., Chalaux, E., Malagelada, J.R., and Real, F.X. (2007). Pancreas transcription factor 1alpha expression is regulated in pancreatitis. Eur J Clin Invest 37, 791–801.

Murtaugh, L.C. (2014). Pathogenesis of pancreatic cancer: lessons from animal models. Toxicol Pathol 42, 217–228.

Petersen, G.M., Amundadottir, L., Fuchs, C.S., Kraft, P., Stolzenberg-Solomon, R.Z., Jacobs, K.B., Arslan, A.A., Bueno-de-Mesquita, H.B., Gallinger, S., Gross, M., et al. (2010). A genome-wide association study identifies pancreatic cancer susceptibility loci on chromosomes 13q22.1, 1q32.1 and 5p15.33. Nat Genet 42, 224–228.

Rooman, I., and Real, F.X. (2012). Pancreatic ductal adenocarcinoma and acinar cells: a matter of differentiation and development? Gut 61, 449–458.

Roy, N., and Hebrok, M. (2015). Regulation of Cellular Identity in Cancer. Dev Cell 35, 674–684.

Roy, N., Takeuchi, K.K., Ruggeri, J.M., Bailey, P., Chang, D., Li, J., Leonhardt, L., Puri, S., Hoffman, M.T., Gao, S., et al. (2016). PDX1 dynamically regulates pancreatic ductal adenocarcinoma initiation and maintenance. Genes Dev 30, 2669–2683.

Shi, G., Zhu, L., Sun, Y., Bettencourt, R., Damsz, B., Hruban, R.H., and Konieczny, S.F. (2009). Loss of the acinar-restricted transcription factor Mist1 accelerates Kras-induced pancreatic intraepithelial neoplasia. Gastroenterology 136, 1368–1378.

Siegel, R.L., Miller, K.D., and Jemal, A. (2016). Cancer statistics, 2016. CA Cancer J Clin 66, 730.

von Figura, G., Morris, J.P.t., Wright, C.V., and Hebrok, M. (2014). Nr5a2 maintains acinar cell differentiation and constrains oncogenic Kras-mediated pancreatic neoplastic initiation. Gut 63, 656–664.

Willet, S.G., Hale, M.A., Grapin-Botton, A., Magnuson, M.A., MacDonald, R.J., and Wright, C.V. (2014). Dominant and context-specific control of endodermal organ allocation by Ptf1a. Development 141, 4385–4394.

Wolpin, B.M., Rizzato, C., Kraft, P., Kooperberg, C., Petersen, G.M., Wang, Z., Arslan, A.A., Beane-Freeman, L., Bracci, P.M., Buring, J., et al. (2014). Genome-wide association study identifies multiple susceptibility loci for pancreatic cancer. Nat Genet 46, 994–1000.

Yachida, S., Jones, S., Bozic, I., Antal, T., Leary, R., Fu, B., Kamiyama, M., Hruban, R.H., Eshleman, J.R., Nowak, M.A., et al. (2010). Distant metastasis occurs late during the genetic evolution of pancreatic cancer. Nature 467, 1114–1117.

